# Sex chromosome turnover in hybridizing stickleback lineages

**DOI:** 10.1101/2023.11.06.565909

**Authors:** Xueling Yi, Dandan Wang, Kerry Reid, Xueyun Feng, Ari Löytynoja, Juha Merilä

## Abstract

Recent discoveries of sex chromosome diversity across the tree of life have challenged the canonical model of conserved sex chromosome evolution and evoked new theories on labile sex chromosomes that maintain less differentiation and undergo frequent turnover. However, theories of labile sex chromosome evolution lack direct empirical support due to the paucity of case studies demonstrating ongoing sex chromosome turnover in nature. Two divergent lineages (*viz*. WL & EL) of nine-spined sticklebacks (*Pungitius pungitius*) with different sex chromosomes (linkage group [LG] 12 in the EL, unknown in the WL) hybridize in a natural secondary contact zone in the Baltic Sea, providing an opportunity to study ongoing turnover between coexisting sex chromosomes. We first identified an 80 kbp genomic region on LG3 as the sex-determining region (SDR) using whole-genome resequencing data of family crosses. We then verified this region as the SDR in most other WL populations and demonstrated an ongoing sex chromosome turnover in admixed marine populations where the evolutionarily younger and homomorphic LG3 sex chromosomes replace the older and heteromorphic LG12 sex chromosomes. The results provide a rare glimpse of ongoing sex chromosome turnover and indicate possible existence of additional but yet undiscovered sex chromosome diversity in *Pungitius* sticklebacks.

**Teaser:** Evolutionarily young sex chromosomes replace the older ones in the hybrid zone of European *Pungitius* stickleback lineages.

## Introduction

In addition to genetic sex determination, sex chromosomes play an important role in reproductive isolation and speciation ^1,2^. Sex chromosome evolution and diversification are thus crucial processes in biodiversification, and yet these processes remain understudied in many taxa. The canonical model proposes that sex chromosomes evolve from autosomes by gaining the master sex-determining gene (e.g., *Sry* in mammals) in a sex-determining region (SDR) that ceases recombination due to sexually antagonistic selection ^3^. Accumulating repeats and deleterious mutations on the sex-limited chromosome (i.e., Y or W) eventually result in their degeneration and structural heteromorphism, which has been observed in highly conserved sex chromosomes of eutherian mammals, birds, and some insects ^4^. On the other hand, recent studies of reptiles, amphibians, and fish have found highly labile sex chromosomes where different (sometimes unknown) genes may be recruited to trigger the sex-determining development ^5^. In addition, the observation of diverse SDRs and homomorphic sex chromosomes indicates frequent turnovers where sex chromosomes can revert to autosomes before becoming highly diverged or degenerated ^6–8^, which indicates a nonlinear evolutionary model with little impact from sexually antagonistic selection ^3,9–11^. The diversity and evolutionary lability of sex chromosomes across the tree of life ^12^ has thus challenged the canonical model and inspired new theories of sex chromosome evolution3,9,10.

Sex chromosome turnover has been mostly inferred on the basis of sister lineages having different sex chromosomes ^7,8,13^. In theory, the evolutionary process of sex chromosome turnover should involve a temporary stage when different SDRs coexist and either is able to determine sex. For example, an ancestral lineage of Medaka was proposed to have both ancestral and new SDRs, the latter eventually taking over and leading to the contemporary species that experienced a sex chromosome turnover ^5^. However, empirical cases of coexisting sex chromosomes remain rare, partly due to knowledge gaps of the SDR diversity in nature and the difficulty of detecting young homomorphic sex chromosomes ^14^. Recent studies of the Japanese soil frog found hybridization between populations having ZZ/ZW and XX/XY sex systems ^15,16^ but these rare case studies have focussed on homologous sex chromosomes, which is different from the turnover between non-homologous pairs of sex chromosomes where the replaced recombining sex chromosome (X or Z) should revert to an autosome ^3^. Therefore, empirical case studies are needed to demonstrate coexisting non-homologous sex chromosomes in nature and test theoretical mechanisms underlying their turnover such as sexually antagonistic selection ^17^ and selection against deleterious mutation load ^18,19^.

The nine-spined stickleback (*Pungitius pungitius*) is a small cold-adapted fish with a circumpolar distribution in the northern hemisphere. The species took multiple trans-Arctic waves of colonization from North Pacific to Europe, forming divergent western (WL) and eastern (EL) European lineages ^20–24^ that have different sex chromosomes ^25^. The EL and non-European populations all have heteromorphic sex chromosomes (XX/XY) identified as linkage group 12 (LG12) with a large Y proposed to have evolved from a historically introgressed inversion from the closely related *P. sinensis* ^25,26^. However, LG12 is not associated with phenotypic sex in the WL which has unidentified homomorphic sex chromosomes that seem to have evolved more recently ^25^. In addition, WL and EL hybridize in the southeastern North Sea and southern Baltic Sea ^20–22,24,27^, indicating coexistence of different sex chromosomes and a potential sex chromosome turnover.

The goal of this study was twofold. First, we aimed to identify the homomorphic sex chromosomes in the WL using whole-genome resequencing data of family crosses with known phenotypic sex. The homomorphic and evolutionarily young WL sex chromosomes are predicted to have low levels of degeneration and a narrow SDR with limited sequence differentiation ^14^. Second, we aimed to characterize sex chromosome diversity using whole-genome resequencing data of 45 nine-spined stickleback populations across the species distribution range, including those in the natural secondary contact zone. We hypothesize that admixture between WL and EL results in a sex chromosome turnover, which predicts an unequal ratio of coexisting sex chromosomes in admixed populations where the more prevalent pair of sex chromosomes take over the other.

## Results

### Identification of the WL SDR

We identified the WL SDR using four one-generation family crosses of wild-caught individuals with known phenotypic sexes from Belgium (Table S1). Whole-genome resequencing of eight parents and 120 F1 offspring generated a total of 2,894,080,759 QC-passed reads that were all mapped to the ver.7 reference genome of *P. pungitius* ^28^. Using the module ParentCall2 in LepMAP3 ^29^, we identified 698 X-associated markers that were mostly (69.05%) located on the LG3 (1898520-18381100 bp) when assuming male heterogamety, but only 340 Z-associated markers that were relatively evenly distributed across the 21 LGs when assuming female heterogamety (Fig. S1A). These results thus indicate LG3 as the most likely sex chromosome pair for the WL nine-spined sticklebacks which is indicated to have male heterogamety (XX/XY).

We further narrowed down the genomic region that has the largest differentiation between sexes (hereafter the SDR), which should include the sex-determining locus and its linked surrounding region that has ceased recombination. To do this, we genotyped and filtered 3,997,985 SNPs across 21 LGs in the EL ver.7 reference genome and compared the two sexes using estimates in 10 kbp non-overlapping windows on each LG (Fig. S1B). No difference was found in sequencing depths between sexes, consistent with the predicted lack of degeneration in the WL sex chromosomes. The inter-sex Fst within each family was around zero in most windows but consistently elevated towards the end of LG3. Most windows had similar heterozygosity in two sexes except for 20 windows on LG3 with the absolute delta heterozygote percentages > 30% between sexes (Fig. 1A). The excessive male heterozygosity is consistent with the XX/XY system. Additional individual-based analyses using inbreeding coefficient (F) and principal component analysis (PCA) on the continuous windows of interest identified only one region in 17260 to 17340 kbp that fully differentiated males and females (Fig. 1B), despite some minor effects of missing data on PCA plots (Fig. S2A ^30^). Accordingly, we identified this 80 kbp region on LG3 as the SDR of the WL population in Belgium.

**Figure 1.**
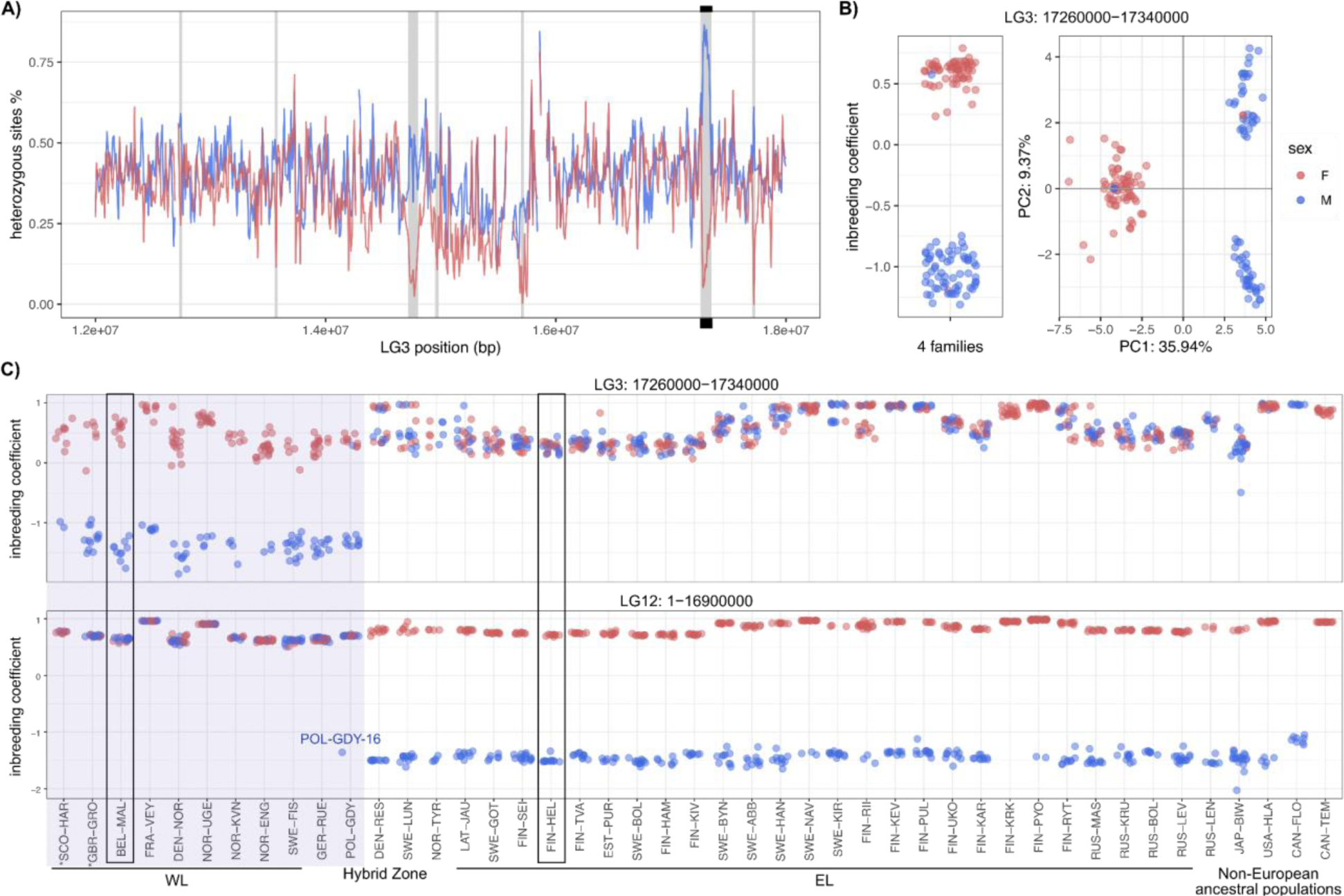
Identification of the LG3 SDR and individual genetic sex. **A)** Window-based estimates of heterozygosity using data of WL family crosses. The x-axis represents the end position of each 10 kbp window. Female values are in red and males in blue (same in the other panels). The grey vertical lines highlight the windows having absolute delta heterozygote percentages > 30% between sexes. Black bars label the position of the identified SDR. **B)** Individual-based analyses (F-values and PCA) of the identified SDR using data of family crosses. Each dot represents one individual and colours represent the phenotypic sex. Two F1 individuals (M15 and F15) from one family cross (Bel_Mal_14) had conflicting genotypic and phenotypic sexes, indicating swapped labels or possibly misidentified phenotypic sex. **C)** The identified genetic sex of 887 wild-caught individuals based on F estimated in the LG3 SDR and the LG12 SDR. Each dot represents one individual and colours represent the identified genetic sex. The 45 populations (Table S2, Fig. 2) are ordered by their indicated sex chromosomes (the 11 LG3-characterized populations are highlighted) and the previously described nuclear ancestry ^24^. The populations in black boxes had known phenotypic sexes and were consistent with the identified genetic sex. Sex identification may not be reliable in the two asterisks-labelled UK populations (see the text).

### Genetic sex of wild-caught individuals and the SDR diversity in European populations

We then characterized genetic sexes and SDRs in all 887 wild-caught nine-spined sticklebacks from 45 globally distributed populations (Table S2) by remapping the published whole-genome resequencing data ^24^ to the EL ver.7 reference genome. A total of 1,447 SNPs were retained in the LG3 SDR (17260-17340 kbp) and 318,850 SNPs were retained in the sex-linked region on LG12 (1-16.9 Mbp, hereafter the LG12 SDR for clarity, but this region probably lacks the most Y-specific SDR and might include some pseudoautosomal fragments ^28^). Genetic sex was identified based on individual inbreeding coefficient (F): low F indicates excessive heterozygosity in the SDR of XY males (similar to the FIS in ^31^), whereas high F indicates excessive homozygosity in the SDR of XX females. In the homologous autosomal regions of each SDR (i.e., LG12 in WL or LG3 in EL individuals), both sexes tend to show high F due to Wahlund effect caused by the joint analysis of all populations. The identified genetic sexes (Fig. 1C) were consistent with known phenotypic sexes available in one WL (BEL-MAL) and one EL population (FIN-HEL, except for one phenotypic female, 16-f, which was also identified as a genetic male in ^24^). Only genetic females were indicated in an EL population (FIN-KRK) but analyses on new samples of known phenotypic males and females showed consistent genetic sex identification (data not shown), indicating biased sampling of this population in our dataset. Overall, most EL populations had the LG12 SDR while WL populations the LG3 SDR, and both SDRs were found in the admixed Polish population (POL-GDY) which had ten LG3-characterized males and one LG12-characterized male (Fig. 1C, Fig. 2).

**Figure 2.**
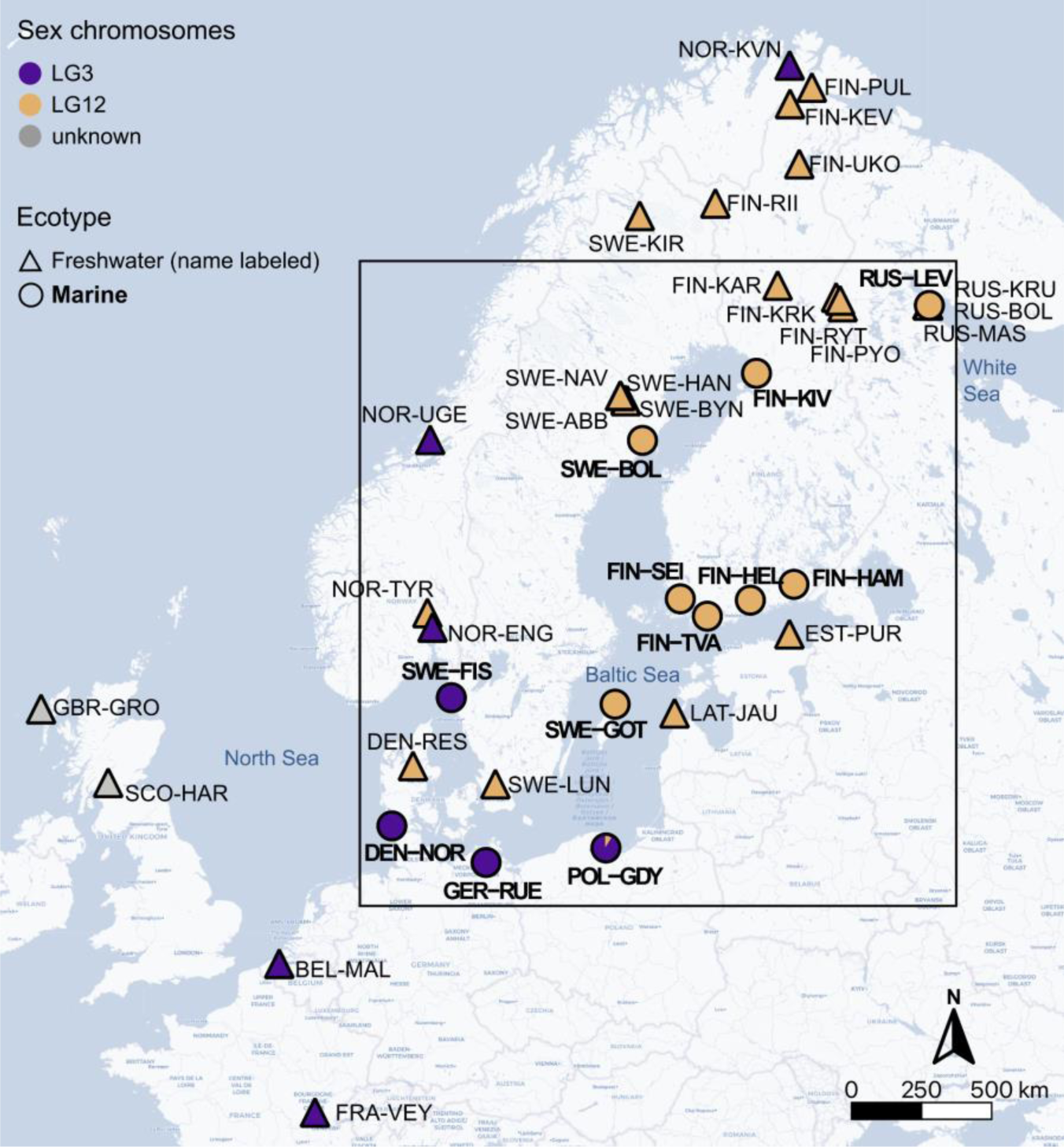
Sex chromosome diversity in the 40 European nine-spined stickleback populations. Each dot represents one population and shapes depict ecotypes. Names of the 12 marine populations are bolded. Colours represent the identified sex chromosomes in each population (both LG3 and LG12 SDRs coexist in a skewed ratio in POL-GDY). The black rectangle highlights the Baltic Sea area.

To further support the genetic sex identification, we conducted PCA on the LG3 SDR using the 11 mostly WL populations (highlighted in Fig. 1C; 221 individuals) and PCA on the LG12 SDR using the 30 mostly EL populations (618 individuals). The ancestral non-European populations were excluded for clarity. No effect of missing data bias was detected and PC1 captured divergence between sexes while PC2 reflected geographic population structure (Fig. S2B,C). The Polish population was included in both PCAs and supported our genetic sex identification of the single LG12-characterized male (POL-GDY-16) and ten LG3-characterized males (Fig. 3A,B). Surprisingly, while PCA of the LG12 SDR in EL showed the expected two clusters of genetic males and females (Fig. 3A), PCA of the LG3 SDR in WL showed three clusters corresponding to two homozygous genotypes and their heterozygous genotype (XY) in the middle (Fig. 3B). This three-stripe PCA pattern has been used as a signature of genomic inversion ^32^. Therefore, we further detected structural variations (SVs) within the LG3 SDR using bam files of paired-end reads mapped to the version 7 reference genome. A putative inversion around 17,275 – 17,328 kbp was detected using males from the family crosses and the LG3-characterized WL populations (Fig. 3F; Table S3). Not all males were detected with this inversion (Table S3), probably due to limitations of next-generation short read sequencing, but no large inversion (>10 kbp) was detected using WL females or individuals from EL or non-European populations (Fig. 3F). Consistently, PCA of the LG3 SDR in the 29 LG12-characterized populations (1,447 SNPs, 598 individuals excluding the Polish population) showed no signal of genomic inversion (Fig. S3). Therefore, these results indicate a putative inversion spanning the majority of the SDR unique to the LG3-Y chromosome.

**Figure 3.**
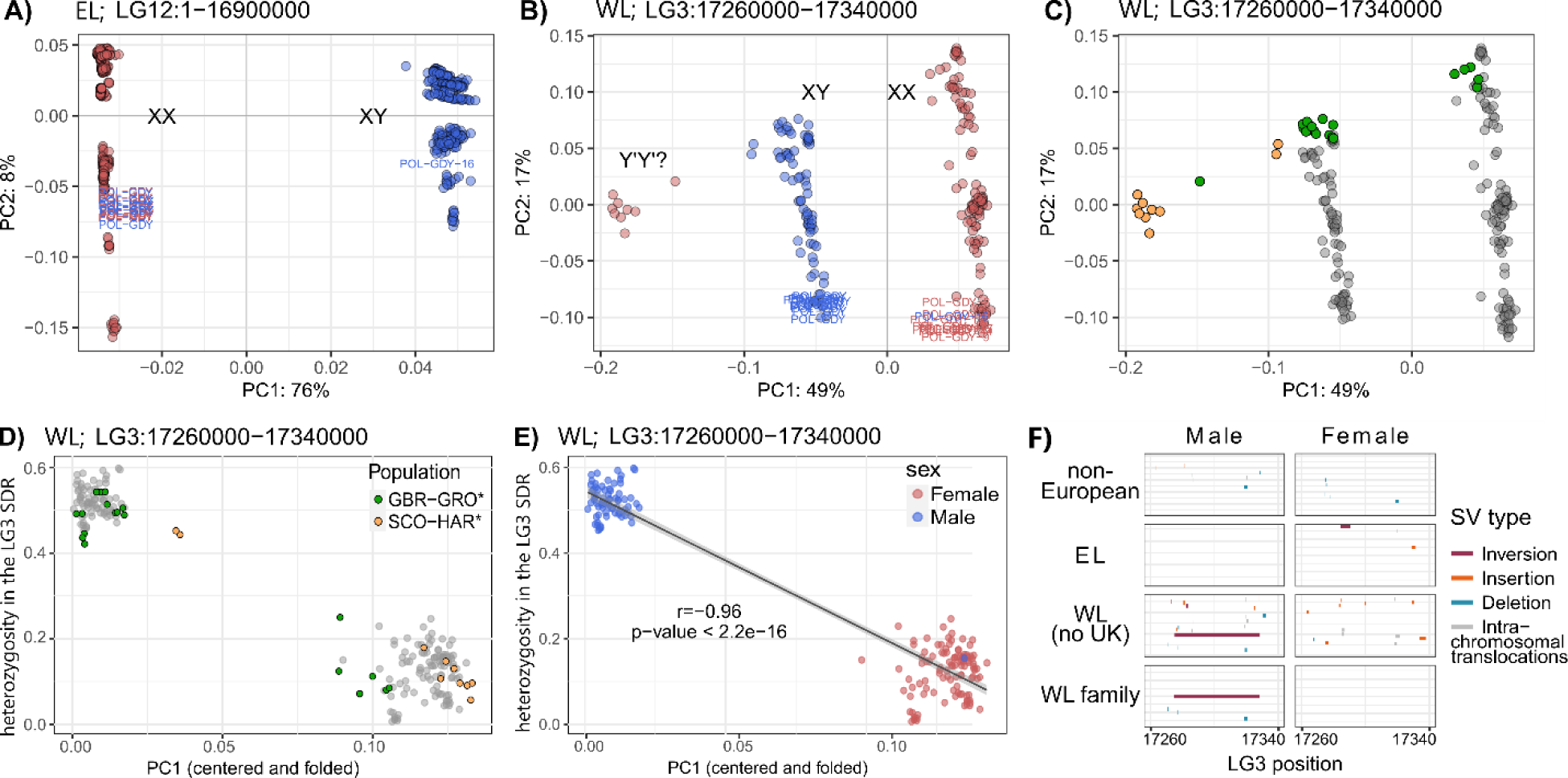
PCA and SV detection indicate a putative inversion in the LG3 SDR. Each dot represents one individual in A-E or detected SV in F, and colours represent the genetic sex (in A, B, E), sampling populations (in C, D), or SV type (in F). **A)** The LG12 SDR of EL populations including POL-GDY (labelled by text). **B & C**) The LG3 SDR of WL populations including POL-GDY. The homozygous Y’Y’ cluster were only found in two UK populations. **D & E)** Individual heterozygosity in the LG3 SDR against the centered and folded PC1 scores from the PCA of LG3 SDR, with D) or without E) the two UK populations. The one homozygous male in the bottom right cluster is the LG12-characterized Polish male (POL-GDY-16). **F)** Putative SVs detected in the LG3 SDR using genetic males or females from different lineages (excluding UK populations, see Table S3). Each bar represents one detected SV colored by type and ordered vertically by the number of supporting reads (high to low from top to bottom).

The PCA plot of the LG3 SDR in WL indicates a homozygous cluster of the inverted YY genotype (Fig. 3B), which is biologically unlikely in such a high frequency even considering rare cases reported in other species, such as mating of sex-reversed individuals ^6^ and viable YY individuals ^7^. In fact, all Y’Y’ (distinct from the sex-related Y haplotype) homozygotes were found in two UK populations which had all three genotype clusters (Fig. 3C). Plots of the observed heterozygosity (percentages of heterozygous sites per individual in the LG3 SDR) against the zero-centered and folded PC1 scores placed the UK individuals around edges or intermediate between the heterozygous and homozygous clusters (Fig. 3D), indicating weaker recombination suppression of this region compared to the other WL populations (Fig. 3E). The UK populations also differ from other WL populations in the SV detection which found one putative inversion shared between the two UK populations and one additional putative inversion unique to GBR-GRO (Table S3). Notably, both inversions were detected in six (out of 13) heterozygous individuals of GBR-GRO (i.e., the middle cluster in Fig. 3C), whereas neither was detected in the five homozygous individuals grouped with WL females (Fig. 3C). These results are intriguing but are limited by the next-generation sequencing data which prevents further interpretations. However, at minimum, these results indicate a different evolutionary history of this LG3 region in the two UK populations which have been also suggested to contain unique historical ancestry ^24^ and strong genetic drift in isolated freshwater habitats. Additional sampling of individuals with known sexes is required to further verify the SDR in these UK populations. Nevertheless, these results indicate potentially still underestimated sex chromosome diversity within this species.

To estimate the origin of this putative inversion(s), we constructed a maximum likelihood (ML) phylogenetic tree using 4,114 SNPs in the LG3 region from nine outgroup individuals and 100 nine-spined sticklebacks including two to three homozygous individuals (i.e., high F in the LG3 SDR; Fig. 1C, Table S2) from each of the 45 populations. The heterozygous individuals (i.e., the XY cluster) were not included because bifurcating trees do not do well with heterozygotes. Our results showed that the LG3-Y’Y’ clade from the UK diverged after the divergence of a basal Western Atlantic lineage (Fig. S4), indicating that the putatively inverted Y’ (and Y) haplotype evolved after the speciation of nine-spined sticklebacks. The LG3-XX clade was grouped within the LG3-autosomal clade composed of EL populations (Fig. S4), consistent with the expected higher similarity between the non-inverted X chromosome and the homologous autosome.

### Hybridization between populations having different SDRs

We next focused on the Baltic Sea hybrid zone ^21,24^ where different SDRs coexist with their autosomal variants. Because freshwater populations are strongly influenced by founder effects and genetic drift, we only analysed marine populations exhibiting more gene flow and larger effective population sizes ^20,33^. A total of 237 individuals were selected from 12 marine populations (low kinship coefficients and relatively equal sex ratio within each population, Table S2), including three LG3-characterized WL populations, eight LG12-characterized EL populations, and the Polish population where both SDRs coexist (Fig. 2). The Danish population (DEN-NOR) from the North Sea and the Russian population from the White Sea (RUS-LEV) were included as representatives of WL- and EL-like ancestors, respectively ^24^. We first characterized the baseline autosomal admixture using 152,546 filtered autosomal SNPs (excluding LG3 and LG12). ADMIXTURE ^34^ indicated optimal four genetic clusters (K=4) based on cross-validation errors (Fig. S5A) and results at K=5 further distinguished the German population (GER-RUE; Fig. 4A). All Polish individuals were highly admixed, including the one LG12-characterized male which is thus not a migrant from an EL population (Fig. 4A). Similar genetic admixture patterns were found when males (n=119) and females (n=118) were analysed separately in ADMIXTURE (Fig. S5B-E), indicating that the prevalence of LG3-characterized males in the highly admixed Polish population could not be explained by higher male-specific gene flow from WL populations.

**Figure 4.**
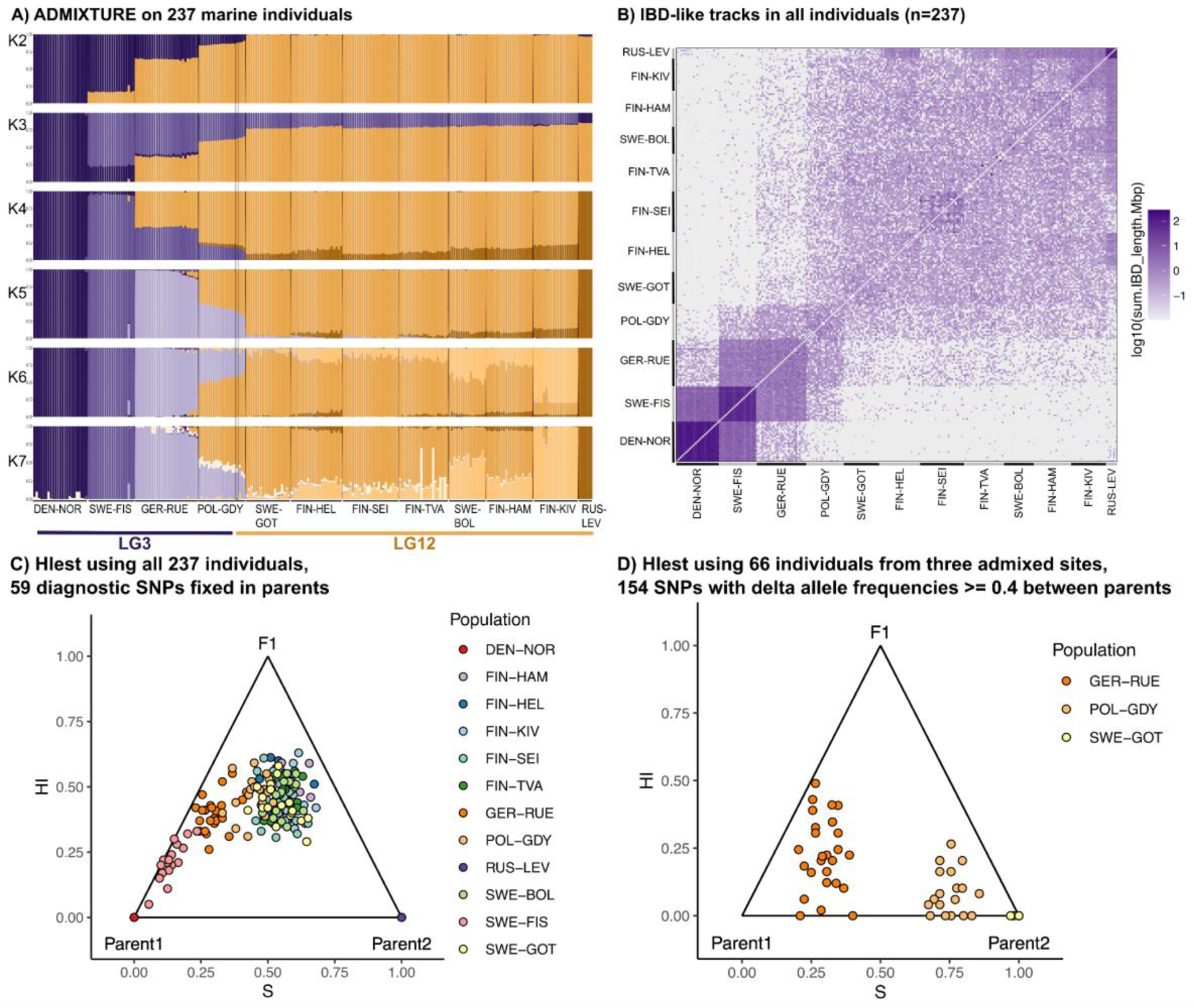
Analyses of the marine populations using the filtered autosomal data. **A)** ADMIXTURE plot of all 237 individuals. The y-axis shows the percentage of inferred ancestry classes indicated by colour. Bars represent individuals ordered by sampling sites and the identified sex chromosomes (labelled at the bottom). The only LG12-characterized male in the Polish population is highlighted. **B)** The total length of IBD-like tracks shared between all pairs of individuals (ordered by population). Darker colors indicate longer shared tracks (in Mbp on the log10 scale). **C & D)** Hybrid classification based on the joint distribution of ancestry (S) and interclass heterozygosity (HI). The expected positions of parental populations and their F1-generation individuals are labelled. Each dot represents one individual and colors depict populations.

To support ADMIXTURE analyses, we estimated fragments of shared ancestry between the 237 marine individuals using identity-by-descent (IBD) tracks. Long tracks represent the exchanged genetic fragments that have not been broken up by recombination and thus relatively recent gene flow, and more shared tracks indicate stronger gene flow between populations ^35^. Total 147,503 IBD-like tracks were detected from the 19 autosomes (Fig. S6A, S7). Track lengths decayed with increasing geographic distances between genetic clusters (Fig. S8), consistent with isolation by distance. Longer tracks were shared within populations and similar patterns were observed when using all individuals combined or separated by sex (Fig. 4B, Fig. S6B), consistent with ADMIXTURE plots and indicating lack of sex-specific patterns. Additionally, the joint estimation of ancestry (i.e., hybrid index) and interclass heterozygosity by HIest classified these admixed individuals as late-generation hybrids (e.g., not F1 or F2 ^36,37^), both when using the Danish and Russian populations as parents (59 fixed diagnostic autosomal SNPs, all 237 individuals; Fig. 4C) and when using the German and Swedish populations as parents (154 autosomal SNPs of absolute delta allele frequencies ≥ 0.4 between parental populations, 66 individuals; Fig. 4D). Therefore, the observed skewed ratio of LG3- and LG12-characterized males in the Polish population cannot reflect putative sex-specific migration during early hybridization, but is the result of evolutionary forces (such as selection) working on several generations of admixture.

### Candidate genes in the LG3 SDR

We scanned annotated genes within the LG3 SDR using the NCBI annotation release of the version 7 reference genome (GCF_902500615.1; search range NW_023616457.1:17260000-17340000). A total of four genes are completely located within this region, including the Olfactomedin-like protein 2B (*OLFML2B*), the E3 ubiquitin-protein ligase RNF31 (*RNF31*), the Potassium voltage-gated channel subfamily B member 2-like (*KCNB2*), and the DnaJ homolog subfamily B member 6a (*DNAJB6A*). In addition, the gene exocyst complex component 3 (*EXOC3*) was partially located in this region. Among these genes, *RNF31* has been found to coregulate steroidogenesis in humans ^40^, and a Z chromosome-linked E3 ubiquitin ligase has been found involved in gonadal differentiation and gametogenesis in the Chinese Tongue Sole ^41^. The DnaJ homolog subfamily B member 13 has been found to participate in spermiogenesis in clownfish ^42^, and proteins in the DnaJ family have been suggested to affect spermatogenesis or sperm function in mammals ^43,44^. Therefore, genes *RNF31* and *DNAJB6A* might also play critical roles in male determination in the WL nine-spined sticklebacks.

## Discussion

Recent studies have demonstrated an incredible diversity of sex chromosomes across the tree of life ^12^ and inspired new theories of labile sex chromosome evolution ^3,9,10^. However, empirical case studies of sex chromosome turnover are rare but are needed to test these new theories of labile sex chromosome evolution. To our knowledge, here we demonstrate the first empirical case of ongoing turnover between non-homologous sex chromosomes in nature. Using the nine-spined stickleback as a model system, we first identified LG3 as the previously unknown homomorphic sex chromosomes of WL and found a narrow 80 kbp SDR, consistent with the expected lack of degeneration and limited sequence differentiation in evolutionarily young sex chromosomes. However, it should be noted that this LG3 SDR was identified using the EL reference genome which represents the autosomal LG3 and may not include the most Y-specific regions that have evolved in the WL. While most of the studied WL populations were identified to carry the LG3 SDR, results from UK populations suggested possible existence of additional, yet unrecognized SDR in WL nine-spined sticklebacks. We hypothesize that the evolutionarily young LG3 SDR evolved in an ancestral lineage that colonized western Europe during an early wave of migration ^24^ and was subsequently spread to other WL and admixed populations by gene flow (Fig. 5A). Moreover, we demonstrated an ongoing sex chromosome turnover in the natural hybrid zone between WL and EL, where the evolutionarily younger homomorphic LG3 SDR takes over the older heteromorphic LG12 SDR, indicating relatively higher fitness of males carrying the LG3 SDR than those carrying the LG12 SDR in the hybrid zone (Fig. 5BC). This observed sex chromosome turnover is consistent with the hypothesis of selection against higher deleterious mutation load on the older non-recombining Y chromosome ^18,19^. Alternatively, this sex chromosome turnover might be driven by natural selection for potential beneficial genes linked with the LG3 SDR. It is likely that multiple selection forces contribute to the observed sex chromosome turnover, but the current data cannot distinguish these alternative hypotheses which need to be tested in future studies. Below we discuss how these findings further our understanding of labile sex chromosome evolution.

**Figure 5.**
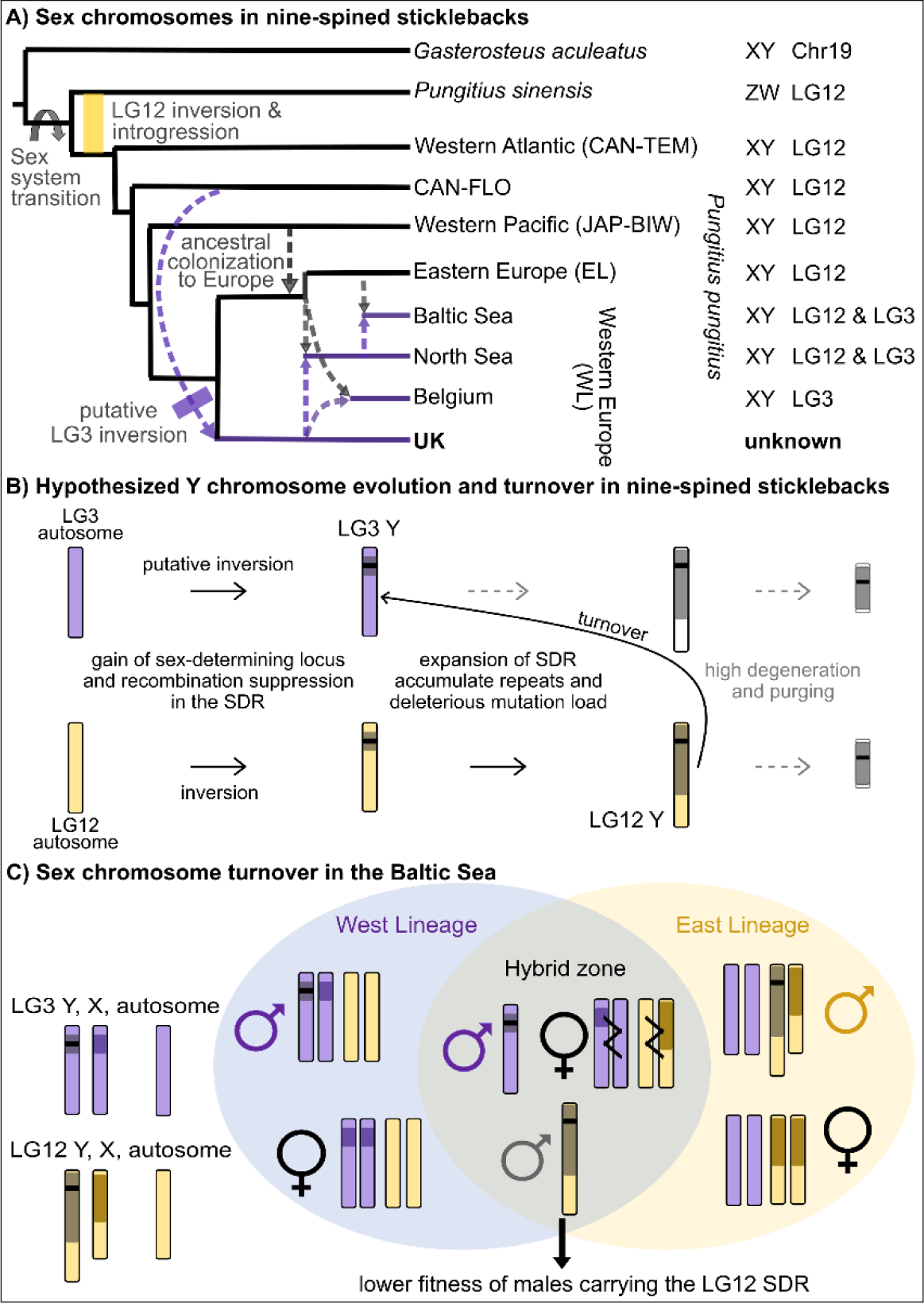
The hypothesized sex chromosome evolution and turnover in nine-spined sticklebacks. **A)** Sex chromosome diversity in nine-spined stickleback lineages. The network is modified from qpGraph plots ^24^. Dashed arrows represent ancestral waves of colonization to Europe and admixture events. The yellow bar represents the previously described LG12 inversion ^25,26^ and the purple bar represents the putative LG3 inversion that might have occurred after an ancestral lineage colonized western Europe. Purple arrows highlight the hypothesized spread of the putative LG3 inversion and possible sex chromosome turnovers. **B)** Hypothesized evolution of the LG3- and LG12-Y chromosomes. Black horizontal bars represent the master sex-determining locus and the surrounding grey regions represent the non-recombining SDR. High Y-degeneration is proposed by the canonical theory but is not found in these labile sex chromosomes. **C)** The proposed sex chromosome turnover in the WL-EL hybrid zone. Dark lines in hybrid females indicate recombination between X chromosomes and their homologous autosomes.

The early evolution of sex chromosomes requires gaining a master sex-determining locus and recombination suppression around this locus in the new SDR. Several master sex-determining genes have been identified in teleost fishes, including *Sox9* ^45^, *Dmy* and *GsdfY* ^46,5^, *Amhr2* ^47,48^, and a duplicated and translocated copy of the anti-Müllerian hormone gene (*AmhY* ^49–52,13^). *AmhY* has been found in several other stickleback lineages including *Culaea* ^13^, the sister genus of *Pungitius* ^22^, but annotations of the nine-spined stickleback reference genome only included the ancestral copy of the *Amh* gene on LG8 ^38^. No master sex-determining gene has been identified so far in any *Pungitius* species which might have recruited different genes to trigger the sex-determining development. The candidate genes identified on the LG3 SDR indicate potential roles in sex determination, but these results are limited by the usage of the EL reference genome which does not contain the true WL sex chromosomes. For the same reason, we were not able to test hypotheses on sexually antagonistic genes (i.e., genes that benefit one sex while harm the other) which have been proposed as the major driving force of recombination suppression of SDR in the canonical model ^53^. However, we hypothesize that the putative inversion of the LG3 SDR might have facilitated the initial recombination suppression (Fig. 5B), similar to previous findings of the EL populations that the LG12 sex chromosomes evolved from an introgressed inversion and that recombination suppression might have predated the sex-determining role of the LG12 SDR ^25,26^. Future studies require high-quality reference assemblies of the WL populations and updated annotations to verify the putative inversion of the LG3 SDR and to better identify candidate sex-determining and sexually antagonistic genes in the WL sex chromosomes.

We demonstrated coexisting SDRs in the natural WL-EL hybrid zone where either LG12-Y or LG3-Y is proposed to be sufficient to determine male development. Because both WL and EL have the XX/XY system, the two Y haplotypes should not co-occur within the same individual as males should not mate with each other (unless there is mating of sex reversed XY females, which has not been reported in this species). On the other hand, females can mate with both types of males and X chromosomes are expected to recombine with their homologous autosomes in hybrid individuals (Fig. 5C), whereas recombination is always suppressed between Y and X or homologous autosomes. The slightly higher EL-like ancestry of the admixed Polish population predicts relatively even proportions of LG3- and LG12-characterized males (or even more of the latter). However, we found only one LG12-characterized male but ten LG3-characterized males, a highly skewed ratio that is unlikely to be caused by sampling effects alone. In addition, our results showed that the observed SDR components are the evolutionary result of multi-generational admixture and cannot be explained by migration of an EL individual, higher male gene flow from WL populations, or sex-specific patterns of introgression. Therefore, we propose that the Polish population has evolved into the condition of having mostly LG3-characterized males from an initial condition of more equal LG3- and LG12-characterized males. If so, this observation might indicate higher fitness of males carrying the LG3-Y compared to males carrying the LG12-Y in the hybrid zone, resulting in a turnover where LG3 replaces LG12 as sex chromosomes (Fig. 5C). Under this scenario, the Polish population probably demonstrates an evolutionary snapshot of ongoing sex chromosome turnover, while the adjacent German population with highly admixed ancestry but only LG3-characterized males might have recently completed this turnover. Similarly, the other freshwater WL populations that currently have the LG3 SDR might also have experienced a sex chromosome turnover, if those locations were initially colonized by EL-like ancestors but later on introgressed by WL-like lineages (such as hypothesized in ^24^).

Sex chromosome turnovers have been proposed to be driven by various mechanisms, including genetic drift ^54^, selection for equal sex ratio ^55,56^, sexually antagonistic selection ^17^, and selection against deleterious mutation load ^18,19^. Genetic drift is unlikely to drive sex chromosome turnover in the admixed marine populations given their large population sizes ^33^. Sex ratio selection is mostly supported in the turnover that includes transitions between XX/XY and ZZ/ZW systems ^16^. Although sexually antagonistic selection has been suggested to drive sex chromosome turnover in some empirical studies ^57,58,8^, recent studies pointed out possibly exaggerated roles of sexual antagonism in the early evolution of sex chromosomes ^11^. Using current annotations, we did not find potential sexually antagonistic genes in the LG3 SDR, indicating that sexually antagonistic selection might not be the driving force of the LG3 SDR evolution or the sex chromosome turnover. However, future studies using the WL reference genome are needed to test whether sexually antagonistic mutations have accumulated on the LG3 sex chromosomes.

The mechanism of selection against deleterious mutation load proposes that when the heterogametic sex is conserved (e.g., XY males in both EL and WL), the evolutionarily younger Y chromosome will take over the older Y chromosome to purge the higher deleterious mutation load accumulated on the non-recombining region of the older Y, while the older X will revert to an autosome to complete the sex chromosome turnover ^18,19^. This mechanism has been suggested to drive frequent sex chromosome turnovers and maintain evolutionarily young sex chromosomes which do not go through major degeneration in some taxa ^3,14^. In nine-spined sticklebacks, the heteromorphic LG12 sex chromosomes are evolutionarily older because they are shared between EL and non-European ancestral populations. Both the longer evolutionary time and the larger SDR range on LG12 sex chromosomes indicate more accumulated deleterious mutations which might have not been purged by degeneration considering the cytogenetically larger Y ^25^. Therefore, the indicated sex chromosome turnover in the WL-EL hybrid zone might be driven by selection against higher deleterious mutation load on the longer and evolutionarily older LG12-Y.

Alternatively, the sex chromosome turnover in the WL-EL hybrid zone might be driven by natural selection favoring the LG3-Y which might contain genes adapted to reproduction in high-salinity environments in the western Baltic Sea ^27^. These fitness-related genes are difficult to identify using the current reference genome containing the autosomal LG3 which is highly different from the LG3-Y (Fig. S4). On the other hand, such genes might also make the LG3-Y less well adapted to the environments of the more eastern Baltic Sea area compared to the LG12-Y, which might explain why the proposed sex chromosome turnover was not observed in the other EL marine populations. However, it is also possible that the LG3-characterized males have not reached these EL populations in high frequency or quantity to initiate the turnover, as nine-spined sticklebacks have low migration rates (e.g., compared to three-spined sticklebacks ^59^). If so, given enough time and continuous gene flow, this sex chromosome turnover might slowly expand to the other EL populations connected to the Baltic Sea, whereas sex chromosomes of the landlocked freshwater populations will likely remain unchanged due to the lack of gene flow (e.g., NOR-TYR and DEN-RES, Fig. 2). Accordingly, the available data cannot distinguish hypotheses of selection against deleterious mutation load on the LG12-Y versus selection for adaptive genes on the LG3-Y, which need to be tested in future studies. It is also likely that these mutually non-exclusive hypotheses both contribute to the WL-EL sex chromosome turnover.

Our empirical study of nine-spined sticklebacks furthers the understanding of labile sex chromosome evolution in several ways. First, we identified the homomorphic sex chromosomes of the WL populations and indicated a putative inversion in the SDR, which might have facilitated the initial recombination suppression. Second, we demonstrated a rare evolutionary snapshot of ongoing sex chromosome turnover in the natural hybrid zone, which is possibly driven by selection against higher deleterious mutation load on the LG12-Y or natural selection favouring putative adaptive genes linked to the LG3-Y. Future studies are needed to test these alternative hypotheses. Lastly, our results showed possibly still unrecognized SDR in the UK populations of nine-spined sticklebacks, indicating high intraspecific sex chromosome diversity which might be maintained by frequent turnovers. Stickleback fishes thus provide a good model system for future studies to further test hypotheses of labile sex chromosome evolution and turnover.

## Materials and methods

### WL family crosses

Four family crosses of the WL nine-spined sticklebacks were generated using four males and four females collected from Brugse polders, Maldegem, Belgium (51°10’ N, 03°28’ E) in 2011 under national and institutional ethical regulations and with permission from the Finnish Food Safety Authority (#1181/0527/2011 and #3701/0460/2011). The paired individuals were artificially mated in a zebrafish rack system (Aquaneering Inc., USA). Fish rearing and crossing followed the descriptions in ^25^. The F1 offspring were reared at 16 °C for 40 weeks maintaining a 24-h photoperiod under license from the Finnish National Animal Experiment Board (#STH379A and #PH1236A). Larvae were fed twice daily with newly hatched brine shrimp nauplii (*Artemia sp*.), and frozen bloodworms (Chironomidae larvae) were added to their diet about eight weeks post-hatching. Individuals were phenotypically sexed based on gonadal inspection. Genomic DNA was extracted from fin clip samples using the salting out method ^60^. Whole-genome resequencing was done by BGI (Hong Kong) using the DNBseq PE 150 platform, targeting 10x for parents and 5x for offspring (15 offspring per sex per family). Demultiplexed raw sequencing reads were processed by AdapterRemoval version 2.3.1 ^61^ to trim adapters, Ns (--trimns), and low-quality reads (--trimqualities). The retained pairs of reads and the collapsed (not truncated) reads were mapped to the ver.7 reference genome ^28^ using bwa-mem in BWA v0.7.17 ^62^ with the -M command and default parameters. The mapped reads were sorted and indexed as bam files and duplicates were marked using SAMtools version 1.16.1 ^63^.

### Identification of the WL SDR

The duplicates-marked reads were first genotyped by mpileup in SAMtools ^64^ using the minimum mapping quality 20 and the minimum base quality 30. Outputs were processed by the module Pileup2Likelihoods of LepMAP3 ^29^. Data mapped to the 21 LGs were processed by the module ParentCall2 to call informative (removeNonInformative=1) markers and identify sex-associated markers that are in non-Mendelian inheritance (diploid in the homogametic sex while haploid in the heterogametic sex) using commands XLimit=2 (assuming XY male heterogamety) or ZLimit=2 (assuming ZW female heterogamety).

Next, the duplicates-marked reads were genotyped by the Genome Analysis Toolkit (GATK) following the best practice protocol ^65,66^. Briefly, raw variants were called using HaplotypeCaller and merged by LG using CombineGVCFs, and all samples were jointly genotyped using GenotypeGVCFs. Biallelic SNPs on each LG were extracted using BCFtools (-m2 -M2 -v snps –min-ac=1 ^67^) and filtered using VCFtools 0.1.16 ^68^ by quality (--minGQ 20 --minQ 30), missingness (--max-missing 0.3), and minor allele frequency (--maf 0.01). The following analyses were conducted in 10 kbp non-overlapping windows on each LG. Mean sequencing depth per site was estimated in VCFtools (--site-mean-depth) using only F1 individuals to avoid bias by sequencing coverage. The ratio of female to male depth was calculated per site, and the average depth ratio across sites was estimated per window. The per-site Weir and Cockerham’s Fst was estimated between F1 males and females of each family cross using VCFtools (-- weir-fst-pop). The percentage of heterozygous sites per sex per window (excluding missing genotypes) was estimated using custom scripts (available on Github). All results were visualized in R v4 ^69^. To verify candidate SDR windows, individual inbreeding coefficient (F) and missingness were estimated in VCFtools and PCA was conducted in the R package adegenet ^70^ using the function glpca with all axes retained.

### Genetic sexes and SDR in wild populations

The published raw whole-genome resequencing data (∼10x) of 887 wild-caught individuals ^24^ were re-mapped to the ver.7 reference genome and processed using GATK in the same way described above. Biallelic SNPs were extracted from the known LG12 SDR (1-16900000 bp ^28^) and the identified LG3 SDR (17260000-17340000 bp), respectively, and further filtered in VCFtools by quality (--minGQ 20 -- minQ 30), depth (--min-meanDP 5 --max-meanDP 25), missingness (--max-missing 0.3), and minor allele frequency (--maf 0.01). Individual F and missingness were estimated in VCFtools. PCA was conducted using PLINK2 (--pca) with the default of keeping the first 10 PCs ^71^.

Structural variants (SVs) in the LG3 SDR were detected by BreakDancer version 1.4.5 ^72,73^ using inputs of bwa-aligned and duplicates-marked bam files mapped to the LG3 of the version 7 reference genome. A configuration file was generated using the script bam2cfg.pl and quality check was done following the protocol ^73^. Read groups were removed if they had the coefficient of variation of the insert size > 0.4 or the percentage of inter-chromosomal read pairs > 4%. The remaining individuals were processed by breakdancer-max to generate a list of putative SVs which were further filtered by confidence score (≥ 99) and the supporting number of reads (> the median of each dataset). BreakDancer analyses were performed separately on genetic males and females in the following datasets: the WL families (all parents and 4 randomly selected offspring per sex per family), the LG3-characterized WL populations (nine populations including POL-GDY but excluding UK, 182 individuals remained after filtering), the LG12-characterized EL populations (28 populations and 575 individuals after filtering), and the LG12-characterized non-European populations (5 populations and 78 individuals after filtering). The two UK populations with unknown sexes (10 individuals from SCO-HAR and 19 from GBR-GRO) were analyzed either combined or separately.

To construct phylogenetic trees of the LG3 SDR, we selected two to three individuals having high F in the LG3 SDR from each of the 45 populations, and included 9 individuals of other stickleback species as outgroups (*Pungitius sinensis*, *P. platygaster*, *P. tymensis*, *P. kaibarae*, and *Gasterosteus aculeatus*). The outgroup samples were collected in previous projects ^22^, sequenced in the same way as in ^24^, and mapped to the ver.7 reference genome and genotyped by GATK as described above. Biallelic SNPs in the LG3 SDR from the total 109 selected individuals were filtered together in VCFtools using the same parameters described above. The filtered data were transformed into phylip format using vcf2phylip ^74^ and the ML tree was constructed in RAxML version 8 ^75^ using the GTRGAMMA model, a rapid search (-f a), and 1000 bootstraps (-# 1000). The best ML tree was rooted by *Gasterosteus aculeatus* and visualized in FigTree (http://tree.bio.ed.ac.uk/software/figtree/).

### Admixture between marine populations in the Baltic Sea

In each of the 12 marine populations, individuals were filtered based on a kinship cut-off of 0.177 (the convention for filtering first-degree relationships) in PLINK2 ^76^. The GATK-genotyped biallelic SNPs of the retained 237 individuals were subset and filtered in VCFtools by quality (--minGQ 20 --minQ 30), mean sequencing depths (--min-meanDP 5 --max-meanDP 25), minor allele frequency (--maf 0.02), and missingness (--max-missing 0.7). Data mapped to the 19 autosomes (excluding LG12 and LG3) were further filtered in PLINK2 by linkage disequilibrium (--indep-pairwise 50 5 0.2), minor allele frequency (--maf 0.05), and missingness (--geno 0.1). ADMIXTURE version 1.3.0 ^34^ was run with 10-fold cross-validation (CV) and ten replicates per K (number of populations) from 1 to 12. The optimal K was indicated by the lowest CV errors. ADMIXTURE outputs were compiled by CLUMPAK ^77^ and visualized using R.

HIest version 2.0 ^36^ was first conducted on all 237 individuals using diagnostic autosomal SNPs that had no missing data and were fixed for different alleles in the parental populations DEN-NOR and RUS-LEV. The function HIest was run with default settings and the data type “allele count”. Next, we focused on the most admixed Polish population using its adjacent populations (GER-RUE and SWE-GOT) as parents. Because stringent filtering for diagnostic SNPs kept fewer than 50 SNPs which would yield low statistical power ^36^, we retained SNPs that had no missing data and delta allele frequencies ≥ 0.4 between parental populations. IBD-like tracks were estimated between all individual pairs using the above filtered data in IBDseq ^78^, with the minimum logarithm of the odds (LOD) score of 4 and all other parameters in default. The correlation between IBD-like track lengths and their LOD scores was visualized as a quality control of the detected tracks.

## Data Availability

Raw sequencing data of the four family crosses are available on NCBI (BioProject PRJNA1023372). The alignment for the phylogenetic analyses is available in supplementary materials. Bioinformatic codes and scripts used in this study are available on Github (https://github.com/xuelingyi/SexChromosome_ninespined_stickleback). The raw sequencing data of wild-caught individuals have been published in ^24^ and can be accessed on the European Nucleotide Archive (ENA) using access code PRJEB39599.

## Acknowledgements

Thanks to Sami Karja, Heini Natri, Takahito Shikano and Yukinori Shimada for their efforts in rearing the family crosses. We also thank the numerous people who provided or helped with sampling of the fish used in analyses. Petri Kemppainen, Paolo Momigliano, and Weixuan Ning are acknowledged for their insightful comments on the PCA and phylogenetic analyses.

## Author Contribution

X.Y., K.R. and J.M. conceived the study. X.F., A.L., and J.M. collected the raw data. X.F. and A.L. processed and remapped the raw sequencing data of wild-caught individuals. X.Y. processed the raw sequencing data of family crosses. X.Y. analysed the processed data with help from D.W. and K.R.. X.Y. visualized results and wrote the first draft of the manuscript. All authors edited the subsequent versions of the manuscript.

