## Supplementary Materials for "Sex chromosome turnover in hybridizing stickleback lineages"

**tree109_LG3sex_a1m3.min4.phy**: The alignment for the phylogenetic analyses on the LG3 SDR region (total 4114 SNPs). The alignment contains 100 individuals of nine-spined sticklebacks (labeled in Table S2) and 9 individuals of the outgroup species *Gasterosteus aculeatus* (sample IDs: NOR-TYR-348, LAT-JAU-15, SWE-GOT-18), *Pungitius kaibarae* (JAP-OMO-26, KOR-HYE-1, RUS-NRI-10), *Pungitius platygaster* (PUN-PLA-G794), *Pungitius sinensis* (PUN-SIN-7a), and *Pungitius tymensis* (PUN-TYM-50a).

**Table S1. Sample information of the family dataset.** A total of 128 individuals (8 parents and 120 $F_1$ offspring) were listed with their sample ID, family ID, and phenotypic sex.

**Table S2. Sample information of the wild-caught individuals.** A total of 887 individuals from 45 populations were sequenced in Feng et al. (2022). The listed genetic sexes and sex chromosomes were identified in this study. The 100 individuals analyzed in the LG3 SDR phylogeny and the 237 individuals analyzed in the marine dataset are labeled.

**Table S3. The BreakDancer-detected putative inversions within the LG3 SDR.** Only the putative inversions larger than 10 kbp are listed. Numbers of individuals and populations refer to those where the corresponding inversion is detected in each dataset. The UK inversion1 was detected in both populations (GBR-GRO and SCO-HAR) whereas the UK inversion2 was only detected in GBR-GRO.

| Dataset | Pos1 | Pos2 | Size (bp) | Ind/total | Pop/total | Classification |
|---|---|---|---|---|---|---|
| **WLfamily_Male** | 17274876 | 17328108 | 52847 | 8/20 | 4/4 families | putative SDR |
| **WL_Male** | 17275034 | 17328533 | 52881 | 22/67 | 8/9 | putative SDR |
| **UK (combined)** | 17275009 | 17328128 | 52894 | 16/29 | 2/2 | UK inversion1 |
| **UK (combined)** | 17275009 | 17328359 | 53135 | 10/29 | 1/2 | UK inversion2 |
| **SCO-HAR** | 17274888 | 17328128 | 52873 | 9/10 | 1/1 | UK inversion 1 |
| **GBR-GRO** | 17275008 | 17328094 | 52964 | 6/19 | 1/1 | UK inversion 1 |
| **GBR-GRO** | 17275008 | 17328359 | 53135 | 10/19 | 1/1 | UK inversion 2 |

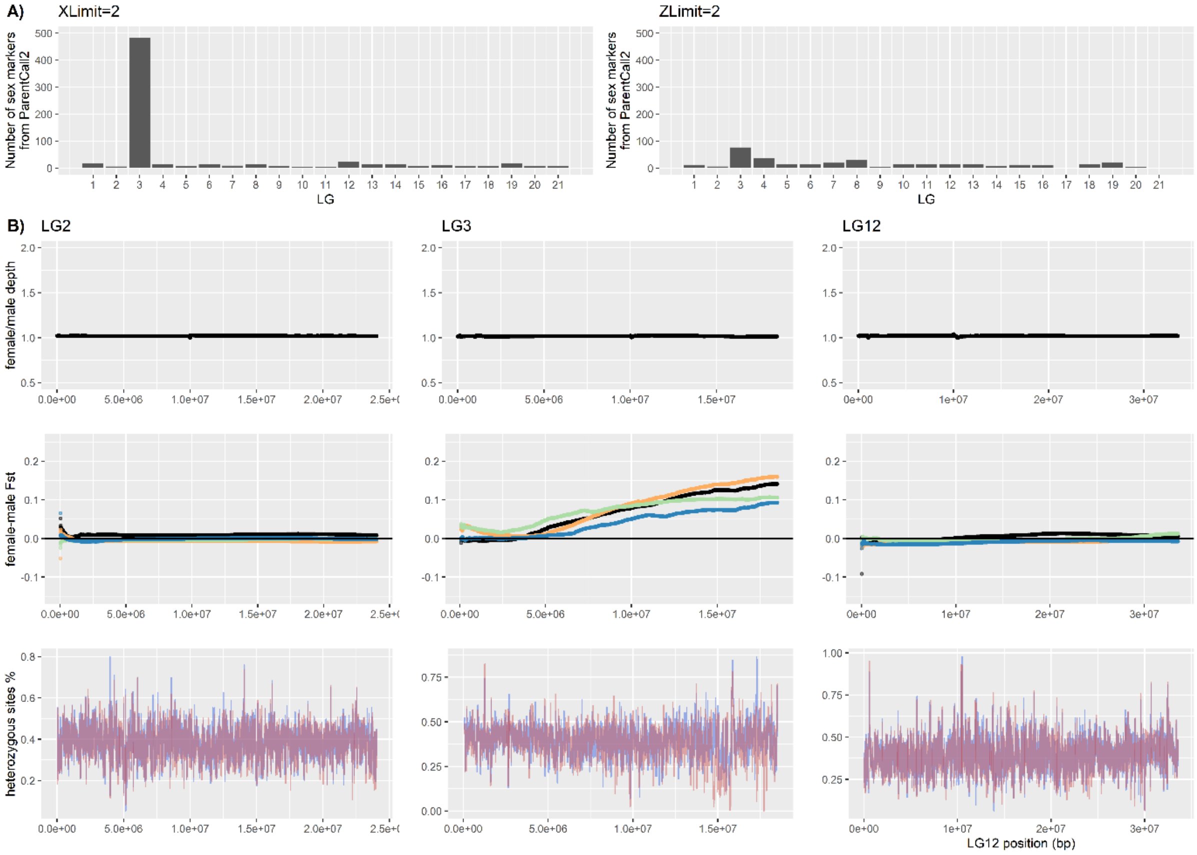


**Figure S1. Identification of the LG3 SDR using data from family crosses. A)** The number of sex-associated markers per linkage group (LG) marked by the ParentCall2 module in LepMAP3 using commands XLimit (left) or ZLimit (right). **B)** Window-based comparisons between sexes in sequencing depth, $F_{st}$ in $F_1$ per family, and percentage of heterozygous sites (top to bottom). Three linkage groups are plotted to represent autosomes (LG2), the identified WL sex chromosomes (LG3), and the known EL sex chromosomes (LG12). The x-axis represents the end positions of each 10 kbp window. The colours of the $F_{st}$ plot represent families: Bel-Mal-11 in black, Bel-Mal-13 in orange, Bel-Mal-14 in green, and Bel-Mal-16 in blue. Horizontal lines represent null expectations of no inter-sex difference ($F_{st}$=0). The colours of the heterozygosity plots represent sex: females in red and males in blue. The deviation between sexes indicates SDR candidates (zoomed in Fig. 1A).

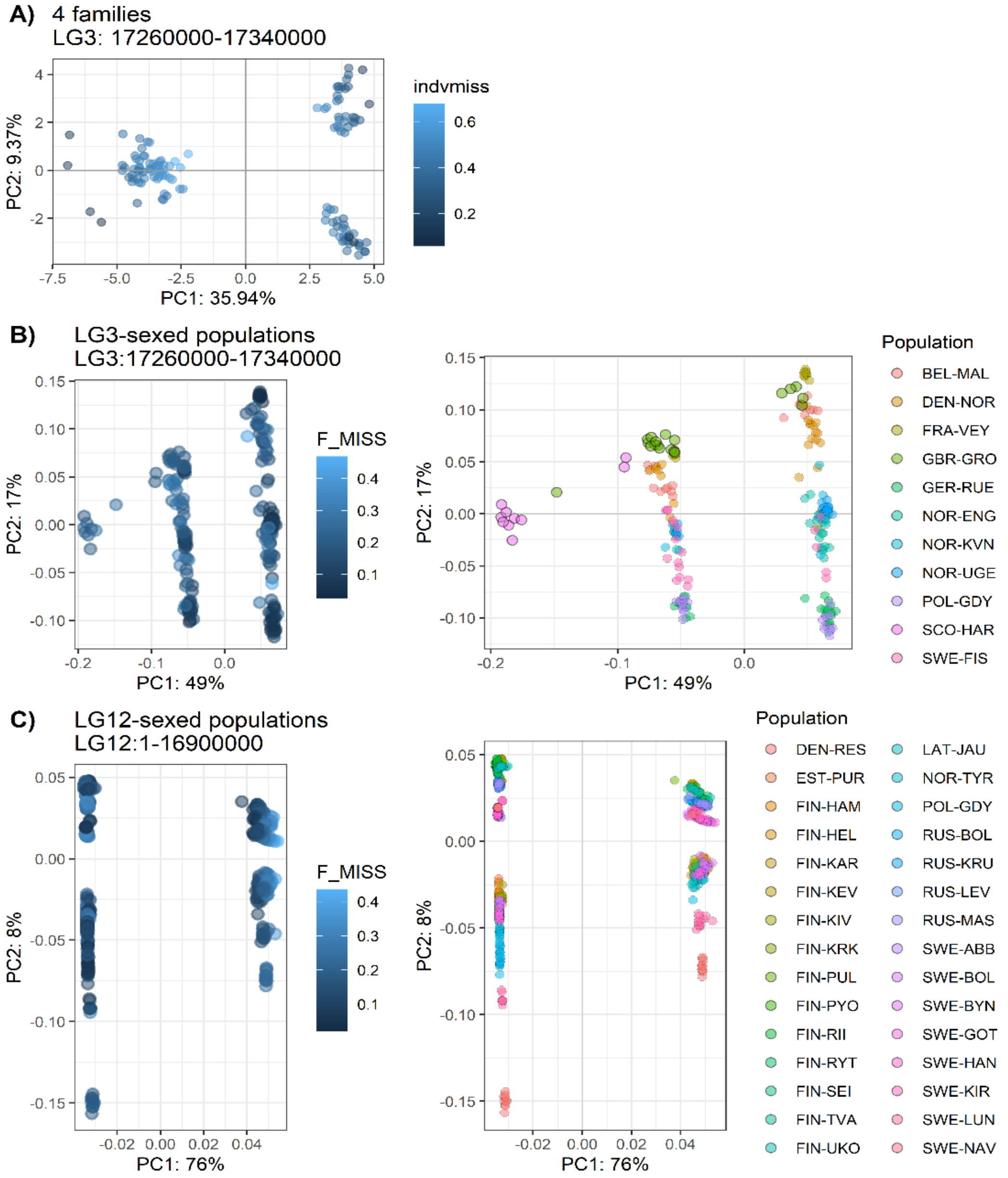


**Figure S2. PCA plots of A) the LG3 SDR in the family crosses, B) the LG3 SDR in WL populations, and C) the LG12 SDR in EL populations.** Each dot represents one individual and is coloured by the proportion of missing data (F_MISS) or population. Individuals in the black outline are from the UK populations (GBR-GRO and SCO-HAR) and might have a different unknown SDR (see text). Individuals having higher percentages of missing data were dragged towards the origin on the PCA plot of family crosses, indicating some extent of bias (Yi & Latch 2022).

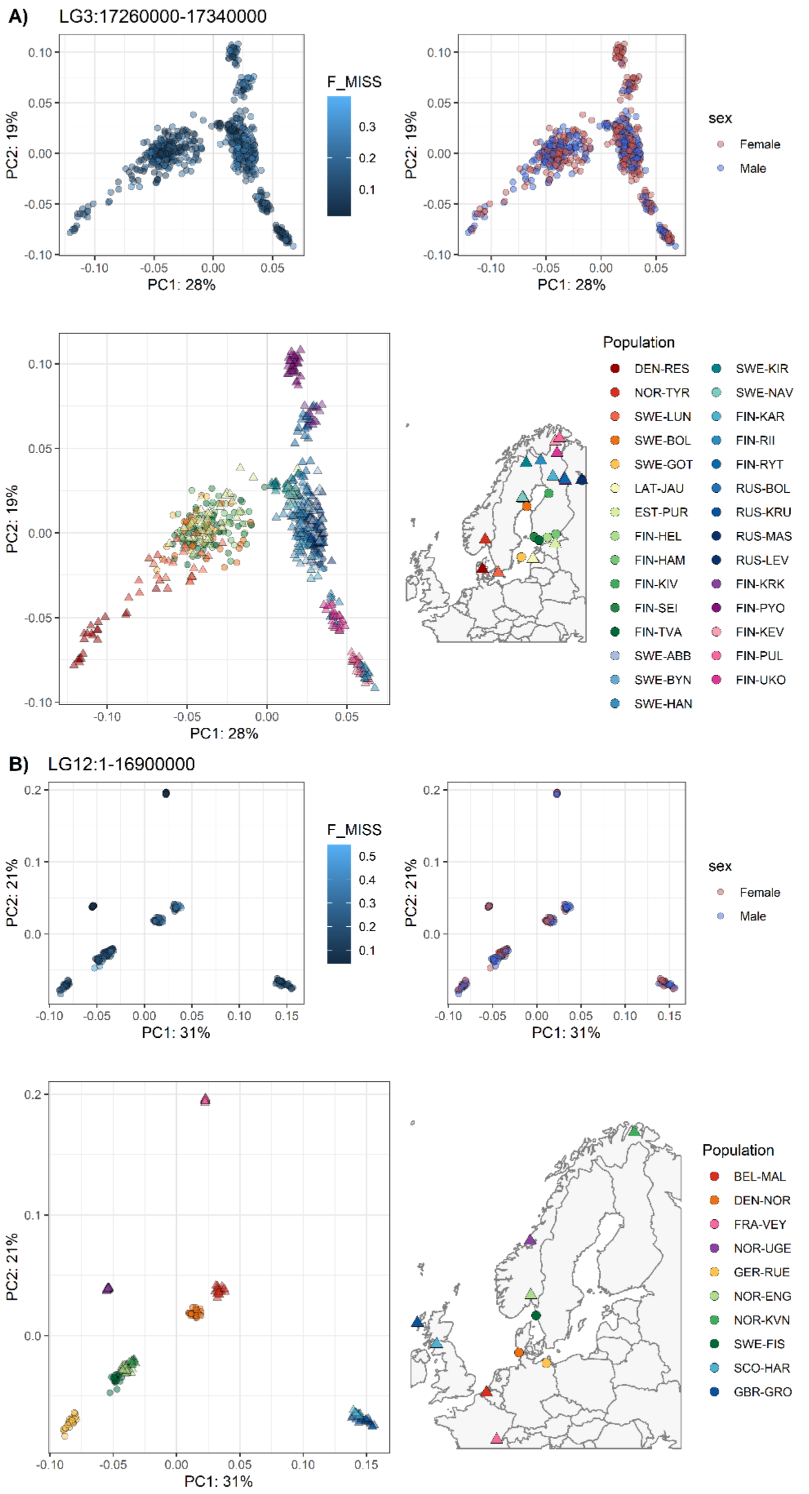


**Figure S3. PCA plots of wild-caught populations and homologous autosomal regions of each SDR.** The population POL-GDY was excluded for clarity. Each dot represents one individual and is coloured by the proportion of missing data (F_MISS), genetic sex, or population. Shapes in maps indicate ecotypes: freshwater in triangles and marine in circles. **A)** The LG3 region in EL populations. PC1 separated populations close to the White Sea from those close to the Baltic Sea and the North Sea. **B)** The LG12 region in WL populations.

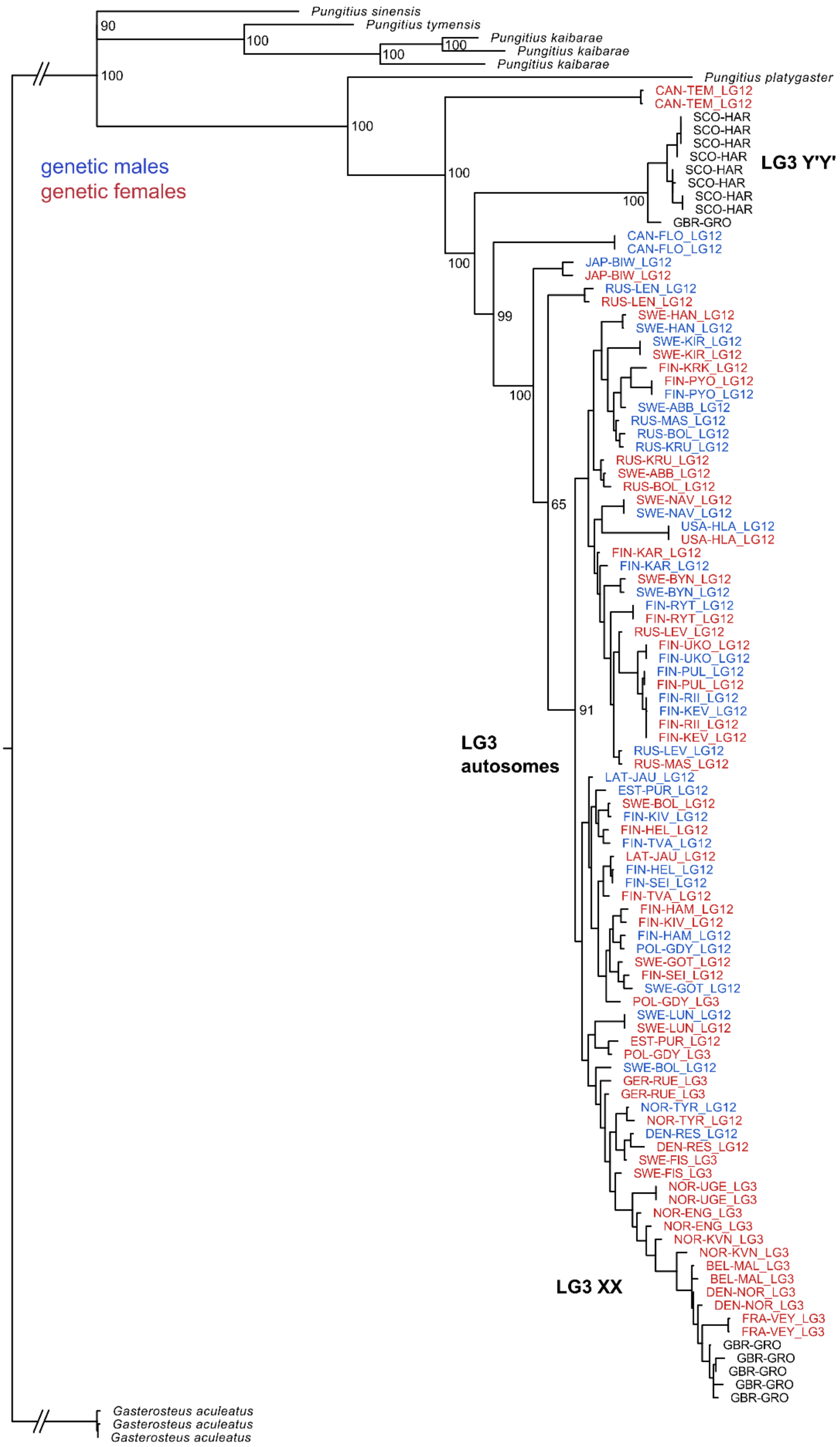


**Figure S4. The maximum likelihood tree of the LG3 SDR.** Bootstrap values of the first few nodes are labelled and outgroup branches are truncated. Each tip represents one individual (only individuals homozygous in this region were included in the analyses). Outgroup tips are labelled by species name in black. Tips of *Pungitius pungitius* are labelled by population and the identified SDR. Genetic females are in red while males are in blue, and the two UK populations with a possibly unknown SDR are in black.

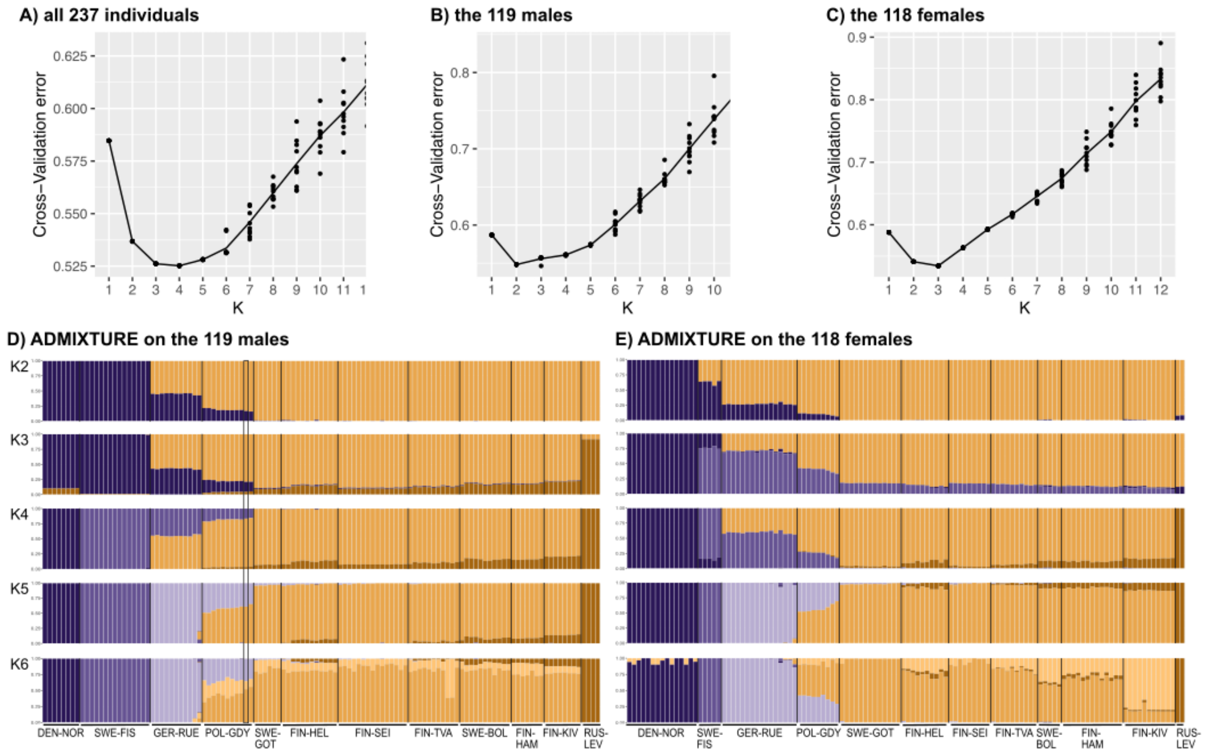


**Figure S5. ADMIXTURE analyses of the marine populations. A-C)** Cross-validation errors using A) the autosomal data of all 237 individuals, B) 119 males, and C) 118 females. Data of 10 replicate runs per K were plotted. The optimal K is indicated by the lowest cross-validation error. **D & E)** ADMIXTURE plots of analyses using D) 119 males and E) 118 females. The y-axis shows the percentage of inferred ancestry classes indicated by colour. Bars represent individuals ordered by sampling sites. The black rectangle in D highlights the only LG12-characterized male in the Polish population.

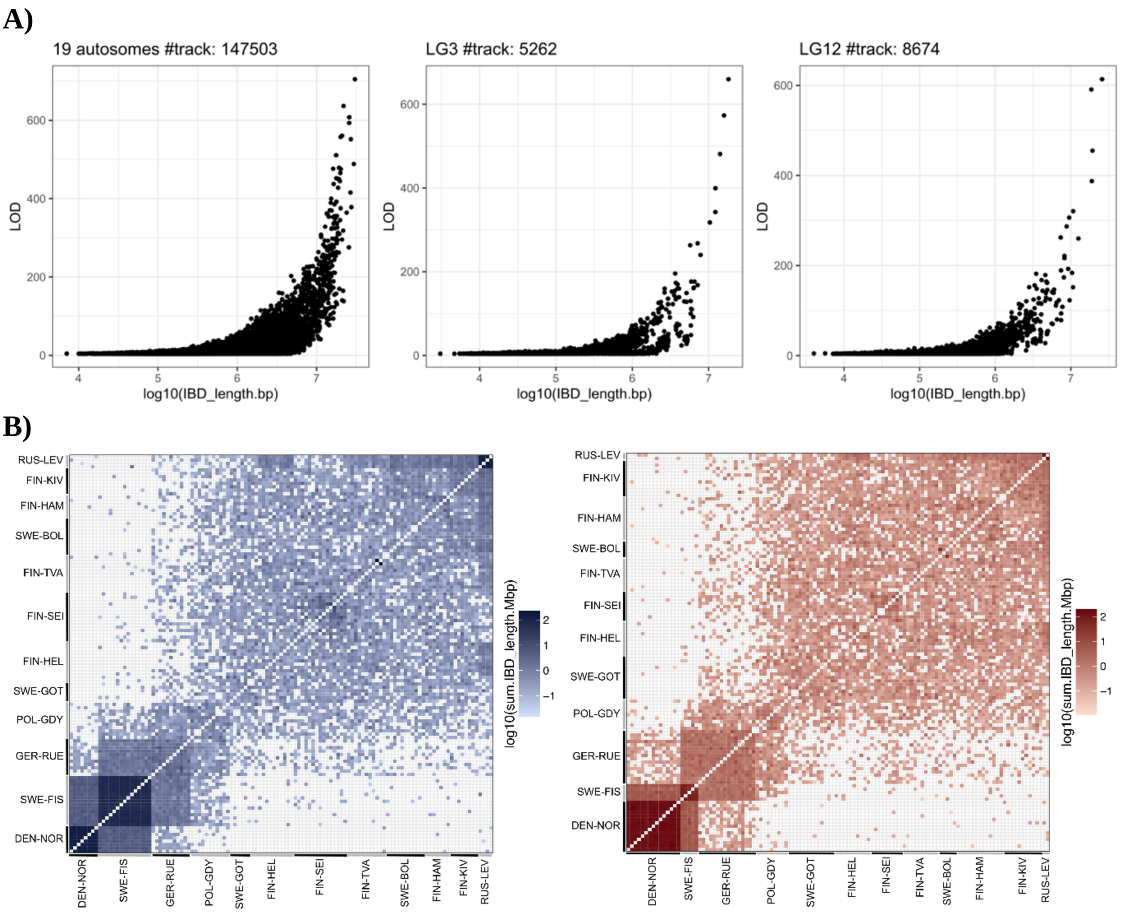


**Figure S6. IBDseq analyses using the marine dataset. A)** Correlation between IBD-like track lengths and LOD scores. Data from the 19 autosomes were combined while data from the sex chromosomes (LG3 and LG12) were plotted separately. All three datasets showed a good correlation between the log10 track length and the LOD score, indicating high quality of the IBD-like tracks (Fontsere et al. 2022). **B)** The total length of autosomal IBD-like tracks shared between pairs of males (left) and pairs of females (right). Individuals are ordered by populations. Darker colors indicate longer shared tracks (in Mbp on the log10 scale).

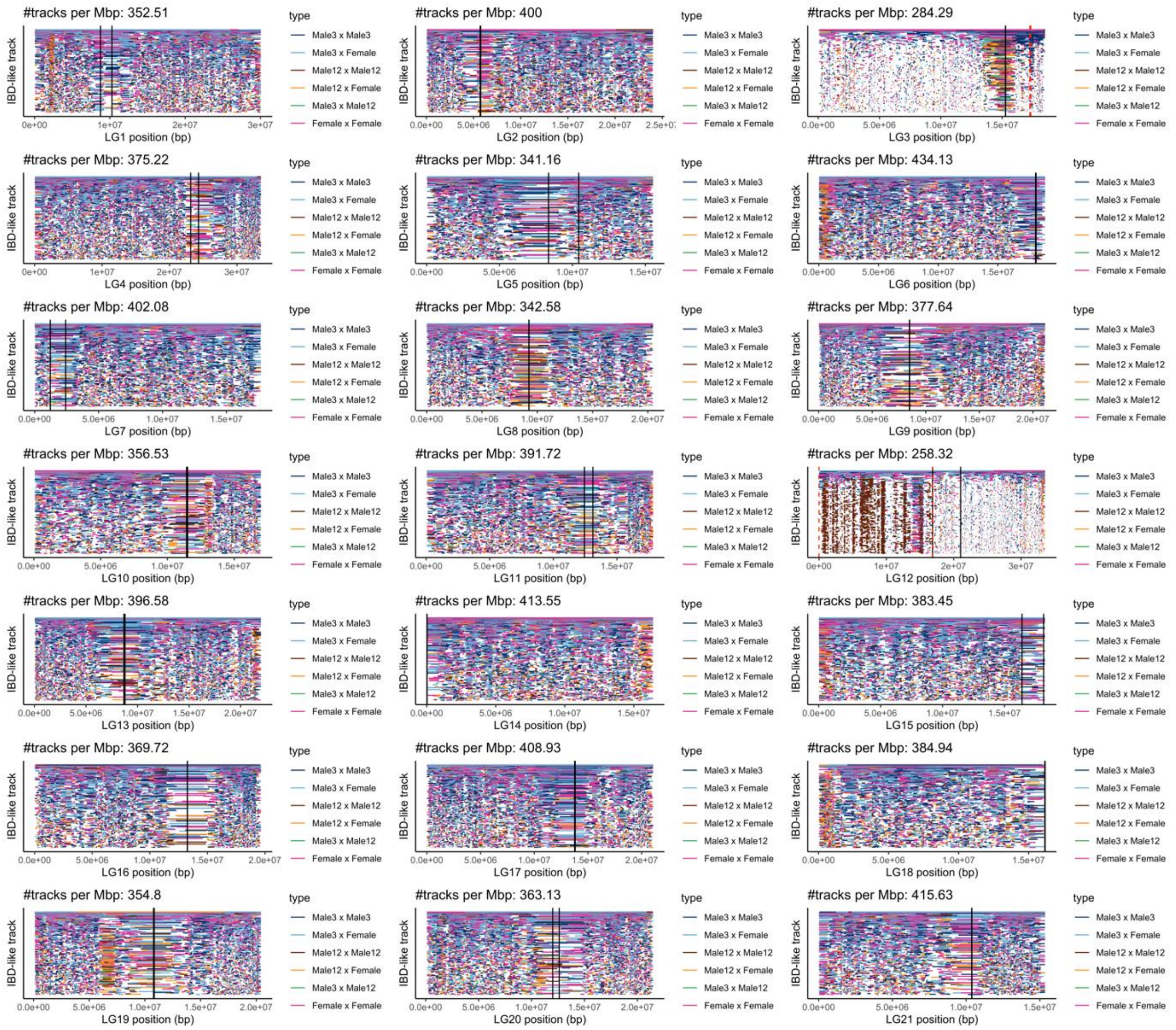


**Figure S7. Distribution of IBD-like tracks per chromosome between individual pairs of marine populations.** Each row is one IBD-like track ordered vertically by LOD scores (low to high from bottom to top) and coloured by the type of individual pair (male3 means LG3-characterized males and male12 means LG12-characterized males). X-axis represents the physical position of each LG. Fewer and shorter tracks were detected on sex chromosomes (LG3 and LG12) whereas the autosomal tracks were relatively evenly distributed among individual pairs and across physical positions. Black vertical lines indicate the start and end positions of centromeric regions in the version 7 reference genome (Kivikoski et al. 2021). The centromeric regions had fewer and longer tracks possibly due to suppressed recombination. Dashed red vertical lines indicate the start and end positions of SDRs (LG3:17260000-17340000; LG12:1-16900000).

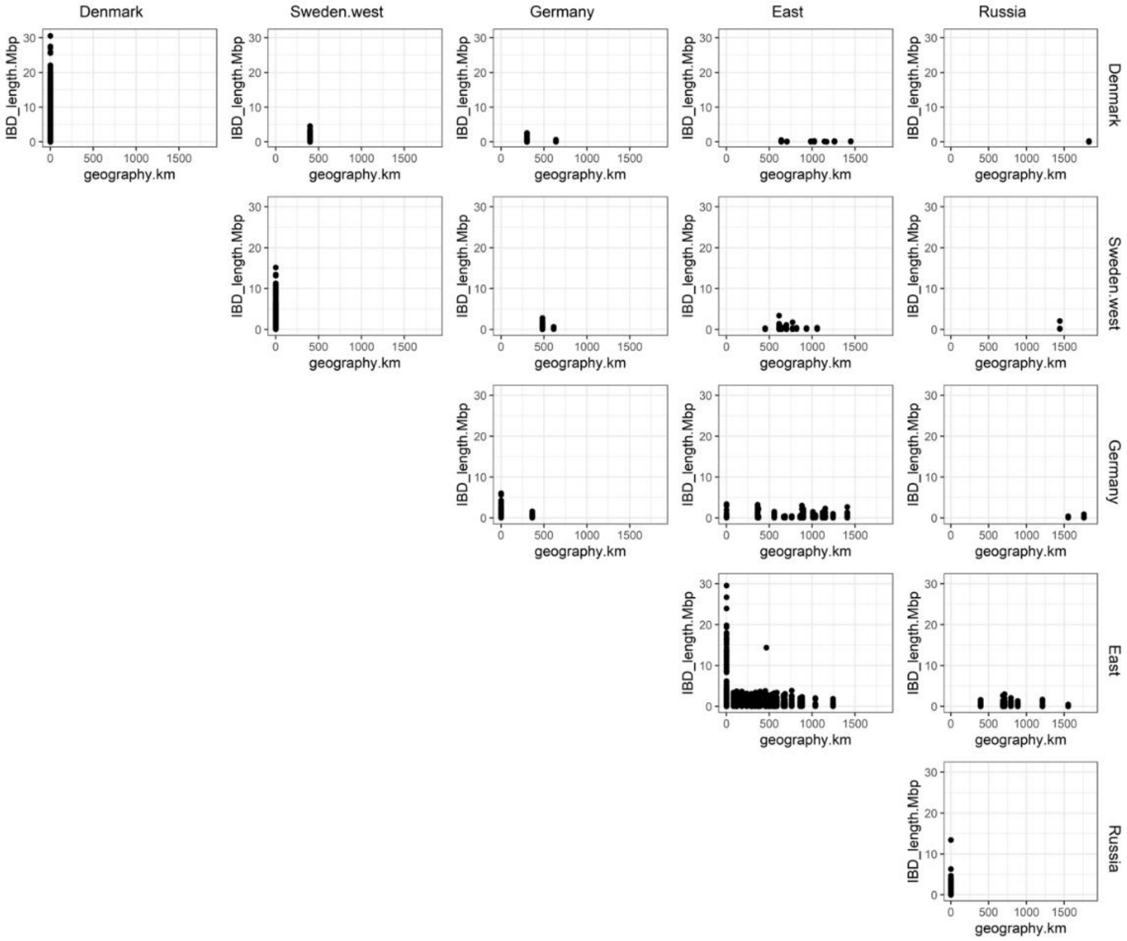


**Figure S8. Correlation between IBD-like track length and geographic distance between genetic clusters using 237 marine individuals.** Assignments to genetic clusters are based on ADMIXTURE results of all individuals at K of 5. The IBD-like tracks detected between distant populations in Denmark and Russia likely represent small parts of shared ancestry from historical introgression (Feng et al. 2022). The outlier IBD-track in the East-East comparison was between a FIN-HEL male (17-m-1) and a SWE-BOL female (SWE-BOL-385).
